# Plasmid-encoded H-NS controls extracellular matrix composition in a modern *Acinetobacter baumannii* urinary isolate

**DOI:** 10.1101/2021.05.19.444899

**Authors:** Saida Benomar, Gisela Di Venanzio, Mario F. Feldman

**Author notes:** Address correspondence to Saida Benomar, or Mario F. Feldman.

## Abstract

*Acinetobacter baumannii* is emerging as a multidrug-resistant (MDR) nosocomial pathogen of increasing threat to human health worldwide. The recent MDR urinary isolate UPAB1 carries the plasmid pAB5, a member of a family of large conjugative plasmids (LCP). LCP encode several antibiotic resistance genes and repress the type VI secretion system (T6SS) to enable their dissemination, employing two TetR transcriptional regulators. Furthermore, pAB5 controls the expression of additional chromosomally encoded genes, impacting UPAB1 virulence. Here we show that a pAB5-encoded H-NS transcriptional regulator represses the synthesis of the exopolysaccharide PNAG and the expression of a previously uncharacterized three-gene cluster that encodes a protein belonging to the CsgG/HfaB family. Members of this protein family are involved in amyloid or polysaccharide formation in other species. Deletion of the CsgG homolog abrogated PNAG production and Cup pili formation, resulting in a subsequent reduction in biofilm formation. Although this gene cluster is widely distributed in Gram-negative bacteria, it remains largely uninvestigated. Our results illustrate the complex cross-talks that take place between plasmids and the chromosomes of their bacterial host, which in this case can contribute to the pathogenesis of *Acinetobacter*.

**IMPORTANCE:** The opportunistic human pathogen *Acinetobacter baumannii* displays the highest reported rates of multidrug resistance among Gram-negative pathogens. Many *A. baumannii* strains carry large conjugative plasmids like pAB5. In recent years, we have witnessed an increase in knowledge about the regulatory cross-talks between plasmids and bacterial chromosomes. Here we show that pAB5 controls the composition of the bacterial extracellular matrix, resulting in a drastic reduction in biofilm formation. The association between biofilm formation, virulence, and antibiotic resistance is well-documented. Therefore, understanding the factors involved in the regulation of biofilm formation in *Acinetobacter* has remarkable therapeutic potential.

## INTRODUCTION

*Acinetobacter baumannii* is regarded as a nosocomial pathogen capable of causing multiple types of infection; however, in the last few years community-acquired infections have become more common. Importantly, *A. baumannii* is a major threat to global health due to the increasing prevalence of the multi-drug resistant (MDR) isolates (1,2). Genes encoding for antibiotic resistance are usually located in chromosomal resistance islands or in plasmids (3,4). Plasmids are major contributors to horizontal gene transfer (HGT), as they facilitate the exchange of genetic material between microorganisms, spreading the MDR phenotype (5-7). For decades, plasmid biology focused on their replication, maintenance, and mobilization, as well as their contribution to antibiotic resistance and virulence (8). More recently, genomic and transcriptomic analyses have begun to uncover complex and dynamic relationships between plasmids and their host chromosome.

*A. baumannii* strains harbor different types of plasmids. Regarding MDR, the Large Conjugative Plasmids (LCPs) family, which are approximately 150-200 Kb, are particularly worrisome. pAB3, the LCP carried in the lab strain ATCC17978, isolated in 1951 carries only one cassette conferring resistance to trimethoprim. However, pAB04 or pAB5, LCPs from recent clinical isolates AB04 and UPAB1, contain 12 and 15 antibiotic resistance cassettes, respectively. This increase in the number of antibiotic resistance cassettes illustrates the rapid evolution of these plasmids. LCPs contain three conserved regions: the MDR region, containing various antibiotic resistance cassettes; a region encoding the T4SS conjugative pilus, required for plasmid dissemination via conjugation; and the regulatory region, which contains several transcriptional regulators (9-11). For example, LCPs harbor two TetR regulators that repress the Type VI secretion system (T6SS) encoded in *A. baumannii* chromosome, which allows conjugation and promotes their own dissemination (10).

We have previously shown that besides providing resistance to antibiotics and repressing T6SS, pAB5 can control the expression of additional chromosomally encoded genes, impacting UPAB1 virulence (11). The regulatory activity of pAB5 can be observed by plating cells with or without this plasmid in Congo-red containing plates. The bacteria colonies carrying pAB5 are much lighter, which reflects a clear reduction in Congo-red binding. Transcriptomic and proteomic analysis revealed that pAB5 reduced the expression of cell-surface components, including (Chaperone/Usher Pathway) CUP pili, β-1 → 6-linked poly-N-acetyl glucosamine (PNAG), and many additional proteins of unknown functions (11). A bioinformatic analysis of the pAB5 sequence revealed that this LCP encodes at least six genes predicted to function as transcriptional regulators. In this work, we explored the cross-talk between pAB5 and the chromosome of UPAB1, and identified the transcriptional regulator involved in regulation of PNAG synthesis. Furthermore, our analysis revealed a novel uncharacterized gene cluster regulated by pAB5, evolutionarily related to curli formation, and whose disruption has a dramatic effect on the surface composition of UPAB1.

## RESULTS

### The transcriptional regulator H-NS, from pAB5, controls UPAB1 phenotype on Congo red binding

We have recently shown that pAB5 regulates the expression of multiple chromosomally encoded virulence factors in UPAB1 (11). One of the most evident phenotypes controlled by pAB5 is congo-red binding, which has been associated to the presence of either amyloid fibers, such as curli, or polysaccharides in *Escherichia coli* and *baumannii* species (ref). pAB5 encodes at least six putative transcriptional regulators (table S1). Among these, there are two TetR regulators, TetR1 and TetR2, virtually identical to the ones encoded in plasmid pAB3. These two TetR regulators inhibit the assembly of the type VI secretion system (T6SS) machinery, which is employed by multiple *Acinetobacter* strains to compete with other bacteria (9,10). pAB5 carries one additional regulator of unknown function, belonging to the TetR family, TetR3. In addition, pAB5 encodes the global regulator H-NS (histone-like nucleoid structuring). H-NS-like proteins have been shown to be implicated in the facilitation of chromosome evolution through their ability to silence transcription, allowing integration of horizontally transferred genes into bacterial chromosomes (12). Finally, orthologues of other regulators, such a FrmR, a putative metal/formaldehyde-sensitive transcriptional repressor, and ArsR, a putative repressor belonging to the arsenic-sensitive family transcriptional regulators are also encoded in pAB5. To investigate if any of these regulators’ controls Congo red binding, we cloned them individually in the pVRL2 expression vector (13). The constructs were transformed into UPAB1p-, and the different strains were platted on Congo red plates. As previously reported, UPAB1 displayed a reduced Congo-red binding (white colonies) compared to UPAB1p-(red colonies), which correlated with the quantification of Congo red binding (fig 1A and B). From the six regulators expressed in UPAB1p-, only H-NS changed the color of the colonies and diminished congo-red binding (fig 1A and B). All other strains expressing the remaining five regulators behaved as UPAB1p-, exhibiting similar leveles of Congo red binding. Finally, UPAB1Δ*h-ns*, a strain carrying pAB5 without *h-ns*, showed comparable levels of Congo-red binding to UPAB1p-(fig 1C and D). These results demonstrate that H-NS is solely responsible for the pAB5-dependent repression of Congo-red binding.

**Figure 1.**
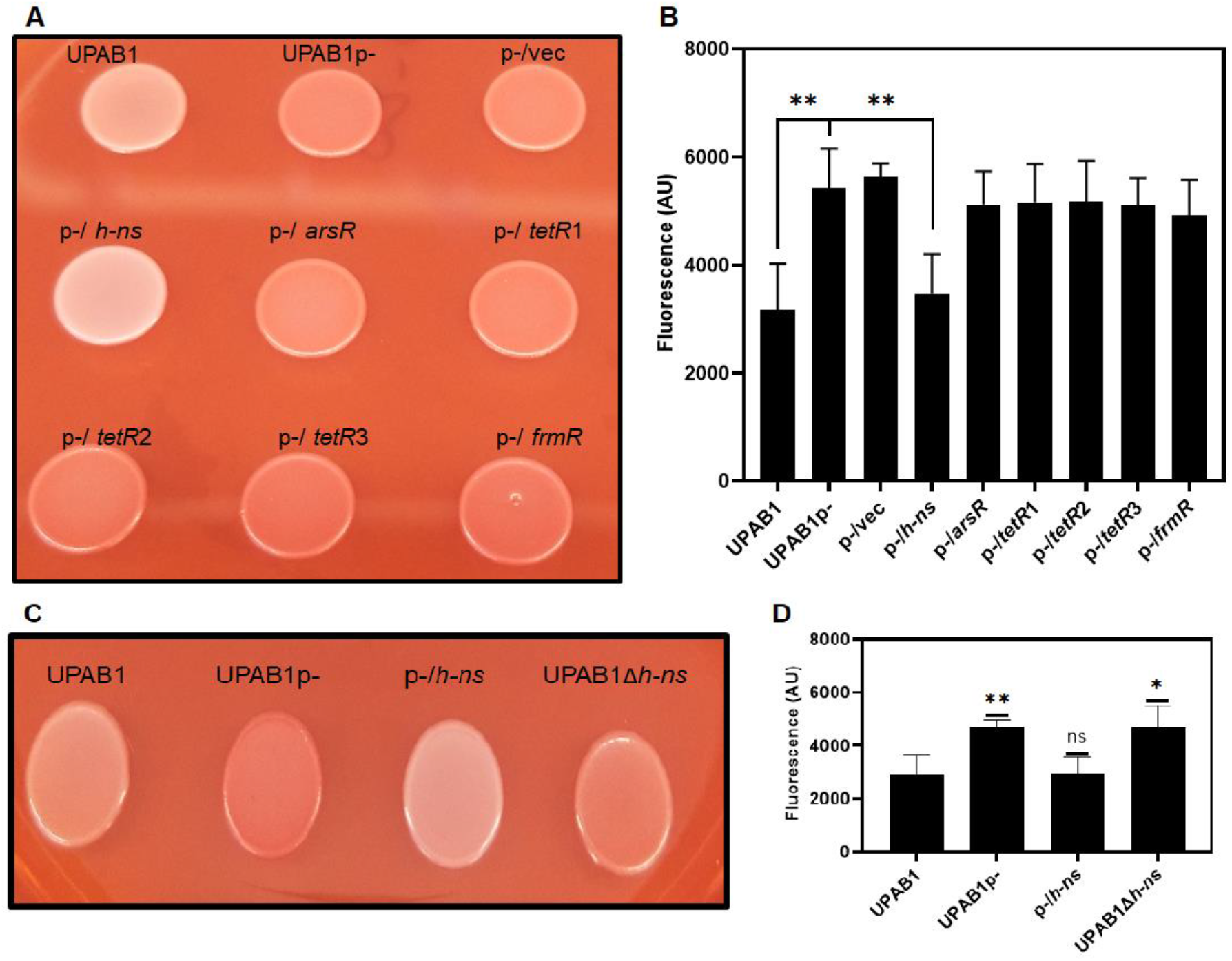
H-NS inhibits Congo red binding. (A & C) UPAB1 and derivatives strain spotted in YESCA Congo-red agar plates. (B & D) Quantification of Congo-red binding (excitation wavelength at 485 nm and the emission at 612nm). The values represent the means and standard deviations from five (B) and four (D) independent experiments. *t* test was performed by comparison with wild type (^**^p≤0.003, ^*^ p≤0.03).

### Plasmid-encoded H-NS inhibits PNAG production

It has been proposed that in some *A. baumannii* strains, Congo-red binding is linked to PNAG production (14). To examine if pAB5 inhibits PNAG production, we used a specific antibody to check for the presence or absence of PNAG. In correlation to congo red phenotypes, UPAB1 and UPAB1p-expressing *h-ns* (p-/*h-ns*) showed drastically reduced levels of PNAG production while UPAB1p-, UPAB1p-harboring the empty vector (p-/vec) and UPAB1Δ*h-ns* strains show similar level of PNAG production (fig 2A). The differences observed on the levels of PNAG production were not related to differences on cells loaded to the membrane (fig 2B). None of the other putative transcriptional regulators from pAB5, altered PNAG-production (fig S1). The locus containing the *pgaABCD* genes has been shown to be responsible for PNAG production in the clinical isolate *A. baumannii* S1 (14). UPAB1 harbors a similar gene cluster containing four genes *pgaABCD* (fig 3A). The pgaA-D cluster encodes for the poly-beta-1,6 N-acetyl-D-glucosamine export porin PgaA; the poly-beta-1,6-N-acetyl-D-glucosamine N-deacetylase PgaB, the poly-beta-1,6 N-acetyl-D-glucosamine synthase PgaC, and the poly-beta-1,6-N-acetyl-D-glucosamine biosynthesis protein PgaD (15). These proteins have an identity ranging from 23 to 55 % to the PNAG cluster in *E. coli* and *Yersinia pestis*. Our recent transcriptomic data showed that this locus is repressed by pAB5 (11). By RT-PCR, we show that expression of *pgaABC* was repressed by H-NS (fig 3B). Furthermore, the pgaA mutant or the whole *pgaA-D* mutant abolished Congo red binding (fig 4A and S2) and PNAG production (fig 4 B,C). These phenotypes were recovered in the complemented strains, demonstrating that *pgaA-D* is responsible for PNAG production in UPAB1. Together, these results show that H-NS downregulates the *pgaA-D* cluster with the concomitant repression in PNAG production.

**Figure 2.**
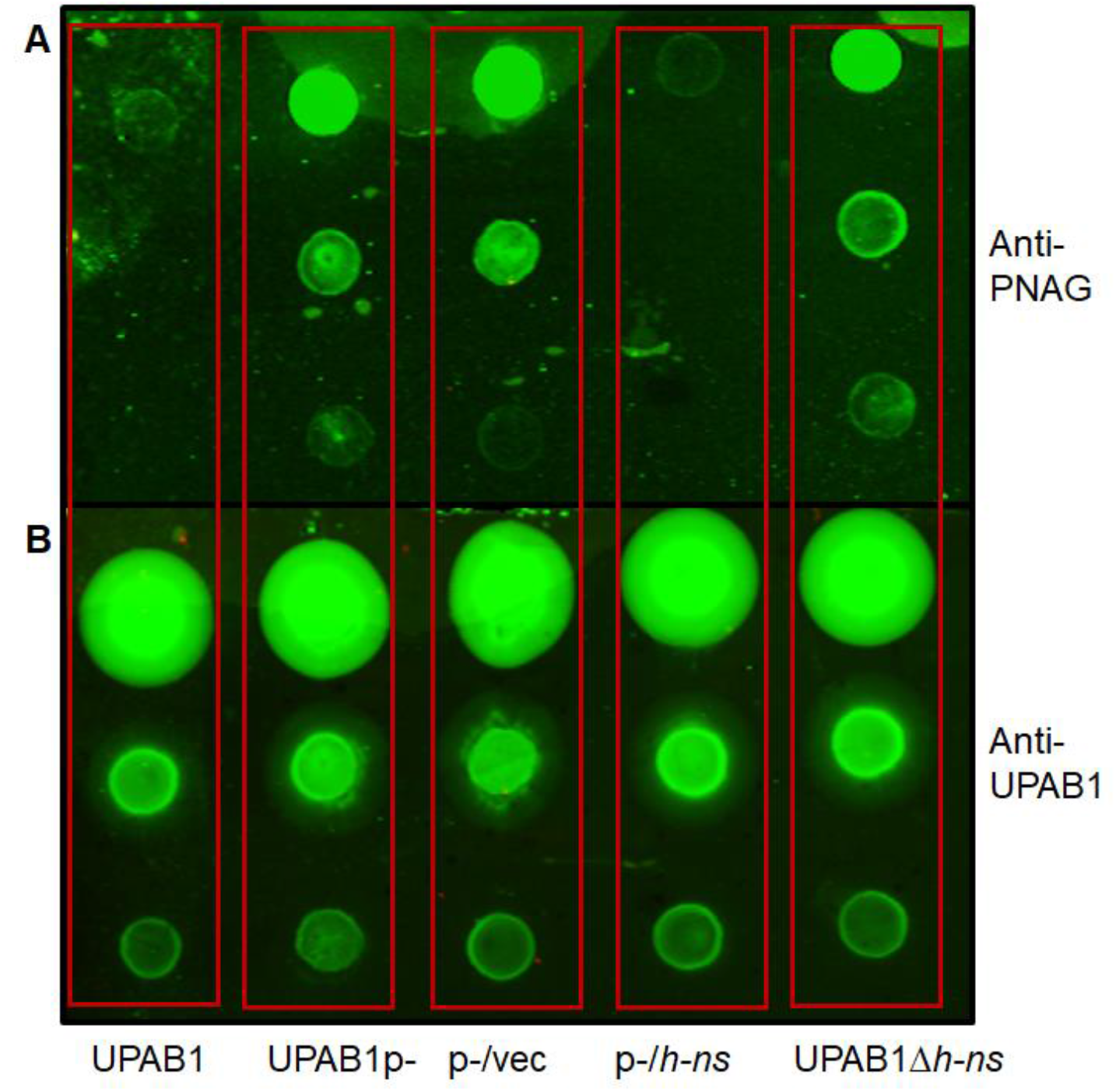
H-NS reduce PNAG production. Immunoblotting using antibodies anti-PNAG (A) and antibodies anti-UPAB1 as a loading control (B). Cells were taken from an overnight LB-agar plate at 26 °C and adjusted to an OD of 1. After treatment, 5 µl of a serial dilution were spotted on nitrocellulose membrane.

**Figure 3.**
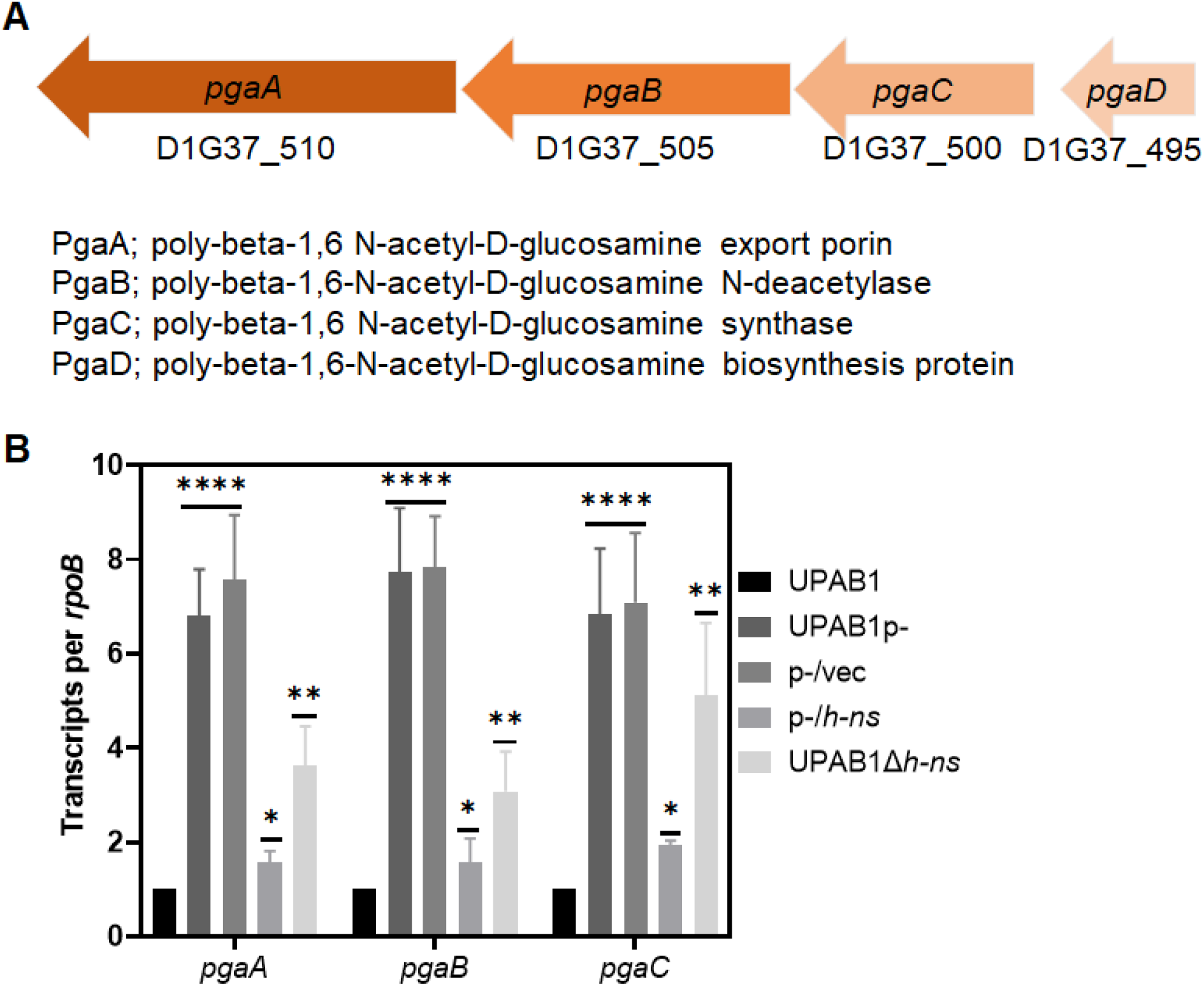
H-NS reduce the expression of *pnag* cluster. (A) Illustration of *pgaA-D* genes cluster. (B) *pgaABC* expression. Transcripts were measured from cells growing on LB plates at 26°C and adjusted to an OD of 1. Results are shown as rpoB-adjusted transcript values. The values represent the means and standard deviations from four independent experiments. T-test (^****^ p≤0.0001, ^**^p≤ 0.005, ^*^ p≤ 0.05).

**Figure 4.**
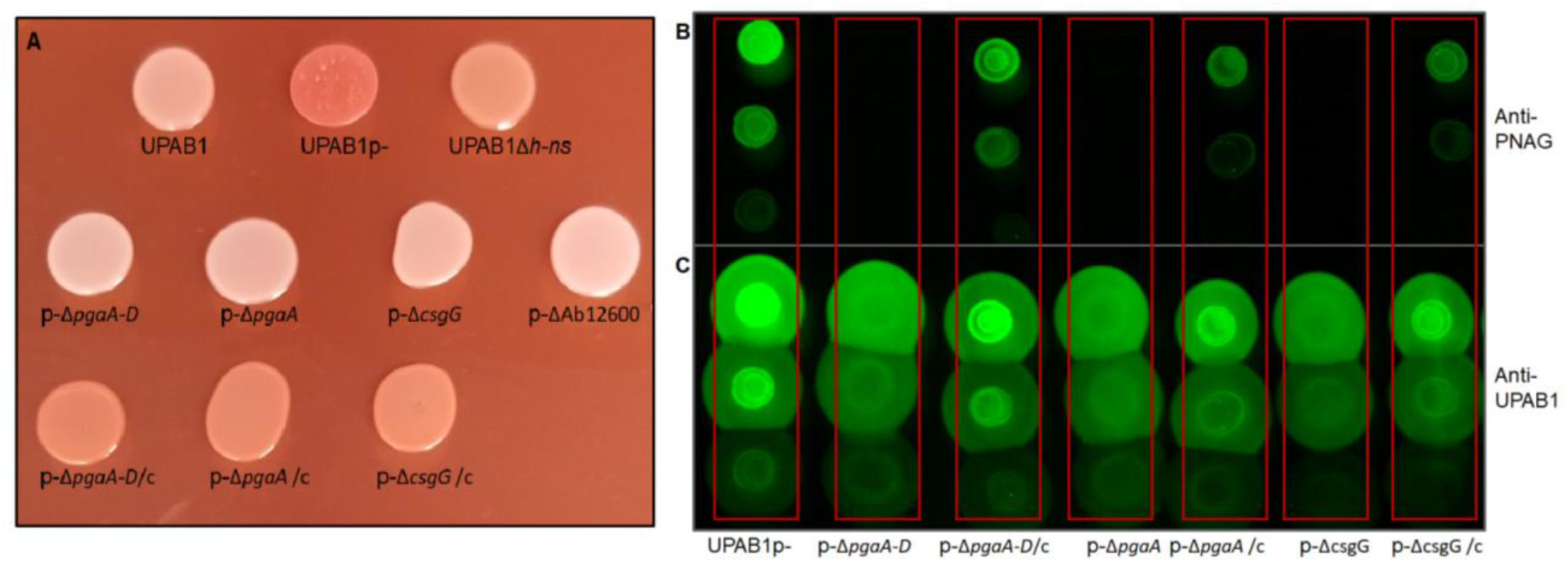
*pgaA-D* and curli-like clusters are involved in PNAG production. (A) phenotype on YESCA-Congo red agar plates of UPAB1p-, derivative mutant and complemented strains. PNAG production was measured using antibodies anti-PNAG (B) and antibodies anti-UPAB1 as loading control (C).

### H-NS regulates the expression of a previously unidentified “curli-like” cluster

Congo-red binding has been routinely employed to monitor Curli amyloid production in *E. coli*. In this bacterium, curli synthesis involves two operons, *csgBAC* and *csgDEFG*. These operons are responsible for curli fiber polymerization, stability, transport and assembly (16-18). Particularly, CsgG forms an oligomeric transport complex and is essential for curli assembly. Although curli formation has not been reported in *Acinetobacter* species, a bioinformatic analysis revealed that a CsgG ortholog (D1G37_12595) is contained within a gene cluster that also comprises two additional genes, D1G37_12600 (“Ab12600”) and D1G37_12605 (“Ab12605”), both encoding putative lipoproteins (fig 5A). Our previous transcriptomic and proteomic analysis indicated that these genes are also downregulated by pAB5 (11). We validated these data by RT-PCR and determined that the transcription of *csgG* and Ab12600 is repressed ∼10 fold by pAB5 or a vector expressing *h-ns* (fig 5B), indicating that, besides PNAG, H-NS also represses the csgG curli-like cluster in UPAB1.

**Figure 5.**
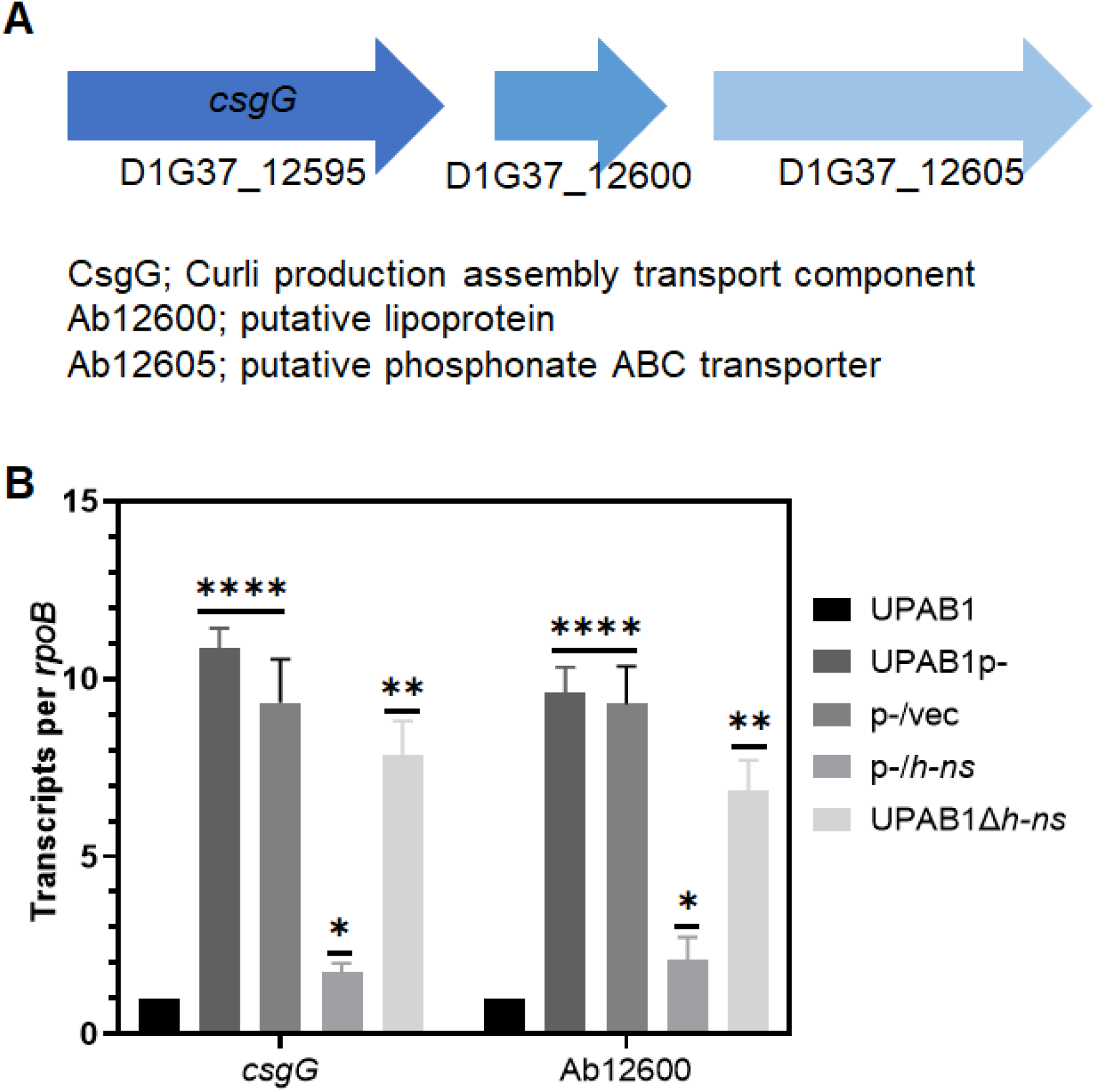
H-NS reduce the expression of curli-like cluster. (A) Illustration of *csgG* cluster. (B) *csgG* and Ab12600 expression. Transcripts were measured from cells growing on LB plates at 26°C and adjusted to an OD of 1. Results are shown as rpoB-adjusted transcript values. Values represent means and standard deviations from four independent experiments. T-test (^****^p≤0.0001, ^**^ p≤ 0.005, ^*^ p≤ 0.05).

### Disruption of CsgG decreases PNAG production, Cup pili formation and biofilm

To explore the role of the CsgG-containing operon in UPAB1, we deleted the *csgG* and 12600 genes in the strains expressing high levels of *csgG*, i.e., UPAB1p- and UPAB1Δ*h-ns*. Surprisingly, *csgG* and 12600 mutant strains showed low levels of Congo-red binding and reduced PNAG production, similar to the *pgaA-D* and *pgaA* mutants, (fig 4A, B and S2). The complementation of the two mutants rescued both phenotypes. To further explore the role of the Curli-like cluster, we analyzed these cells via SEM and TEM. SEM images showed that UPAB1p-cells were coated with a thick layer of extracellular matrix material (fig 6). The *csgG* mutant displayed drastically reduced attachment to the coverslip and, except for some fibers, lacked most of the extracellular matrix (fig 6). This phenotype was partially complemented by expressing the CsgG gene *in trans*. (fig 6). For comparative purposes, we also examined the Δ*pgaA-D* mutant strain. The Δ*CsgG* and Δ*pgaA-D* strains exhibited similar phenotypes, although the reduced binding to the coverslips was less pronounced in the Δ*pgaA-D* strain (fig 6). These phenotypes are not due to growth defects (fig S4), and they correlate with the lower PNAG production in the *csgG* mutant. Our TEM analysis showed that CsgG, but not PgaA-D deletion, results in an almost total abrogation of CUP pili formation (fig 7). CUP pili levels were restored in the complemented cells. Western-blot analysis confirmed that deletion of CsgG abrogated CUP pili expression (Fig S4). Moreover, this analysis confirmed that CUP pili is repressed by pAB5, although this repression was independent of H-NS (Fig S4). The changes in the extracellular matrix were also reflected in the levels of biofilm formation, as the *pgaA-D* and *csgG* mutants produced less biofilm compared to wild-type bacteria (Fig 8). The reduction of biofilm formation was restored in the complemented strains (fig S5). Moreover, cells carrying pAB5, but not pAB5Δ*h-ns*, displayed lower levels of biofilm formation. Together these experiments demonstrate that deletion of CsgG results in reduced production of PNAG and CUP pili, with the subsequent reduction in biofilm formation.

**Figure 6.**
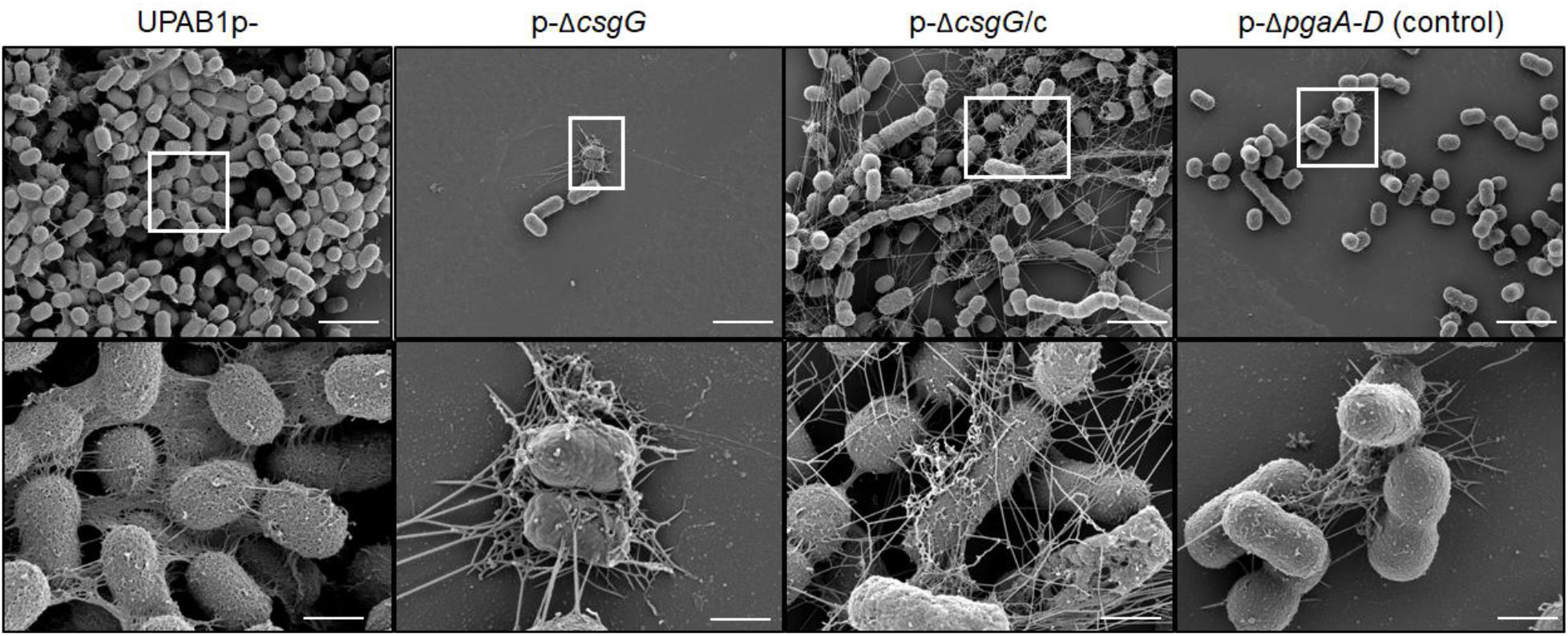
CsgG and PNAG are involved in extracellular matrix production. SEM analysis of UPAB1p-, p-ΔcsgG, p-ΔcsgG/c, and p-ΔpgaA-D (as a control). Cells grown in YESCA media with 4% DMSO in 24-well plate with glass coverslips. After overnight growth, glass coverslips were removed, washed (150 mM cacodylate buffer with 2mM CaCl2), fixed (2.5% glutaraldehyde, 2% paraformaldehyde, and 0.2% tannic acid in 150 mM cacodylate buffer (pH 7.4) with 2mM CaCl2) and treated for observation. The bottom panel is a magnification of the white square in the top panel. Scale bars are 1 μm for top panel and 200 nm for bottom panel.

**Figure 7.**
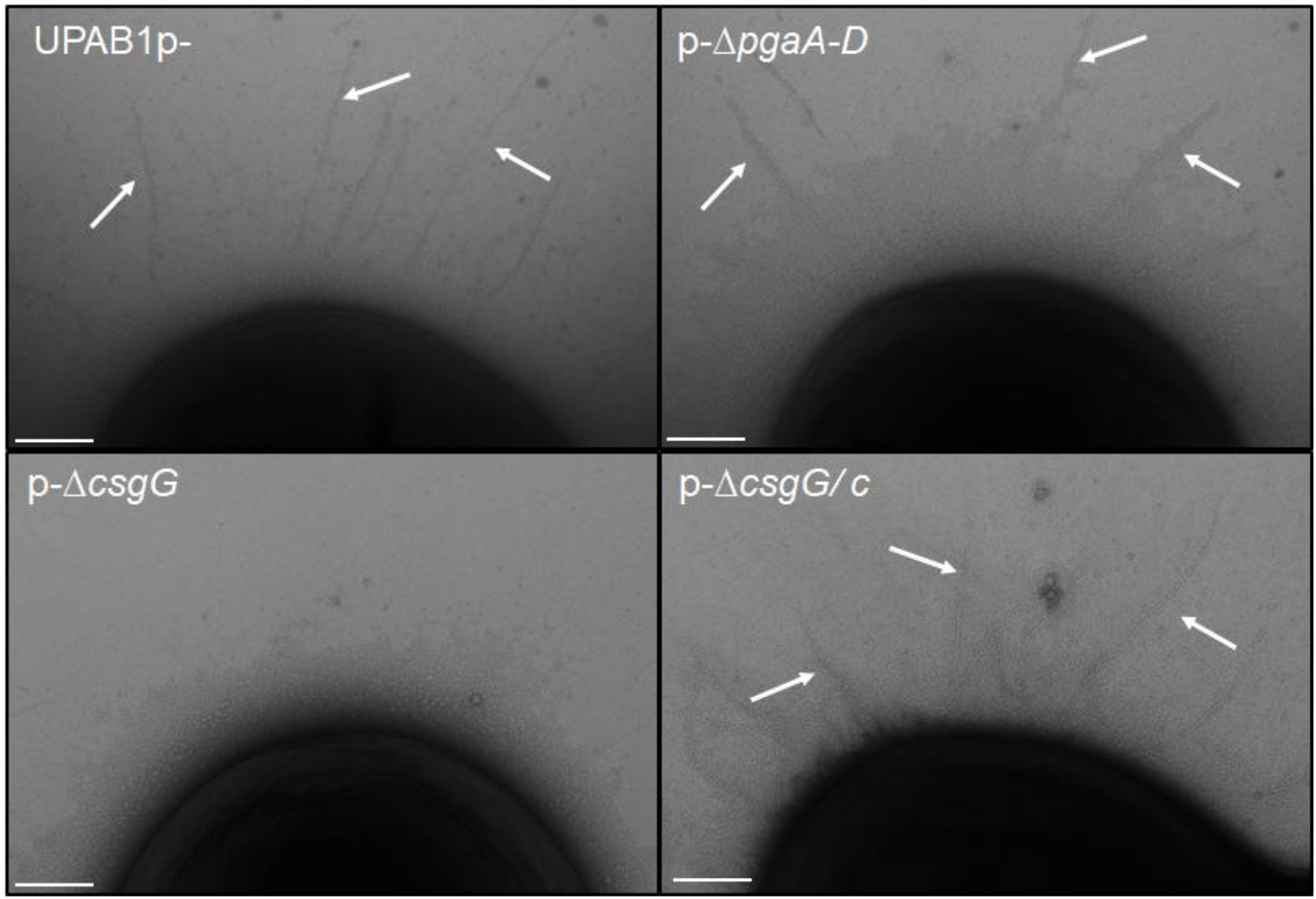
CsgG is involved in CUP pili formation. Transmission electron microscopic images showing CUP pili. The pili structures were absent in p-Δ*csgG* strain and restored in the complemented strain (p-Δ*csgG*/c). Cells grow for 48 hours in YESCA media supplemented with 4% DMSO with shaking at 26 °C. Sclae bar 100 nm

**Figure 8.**
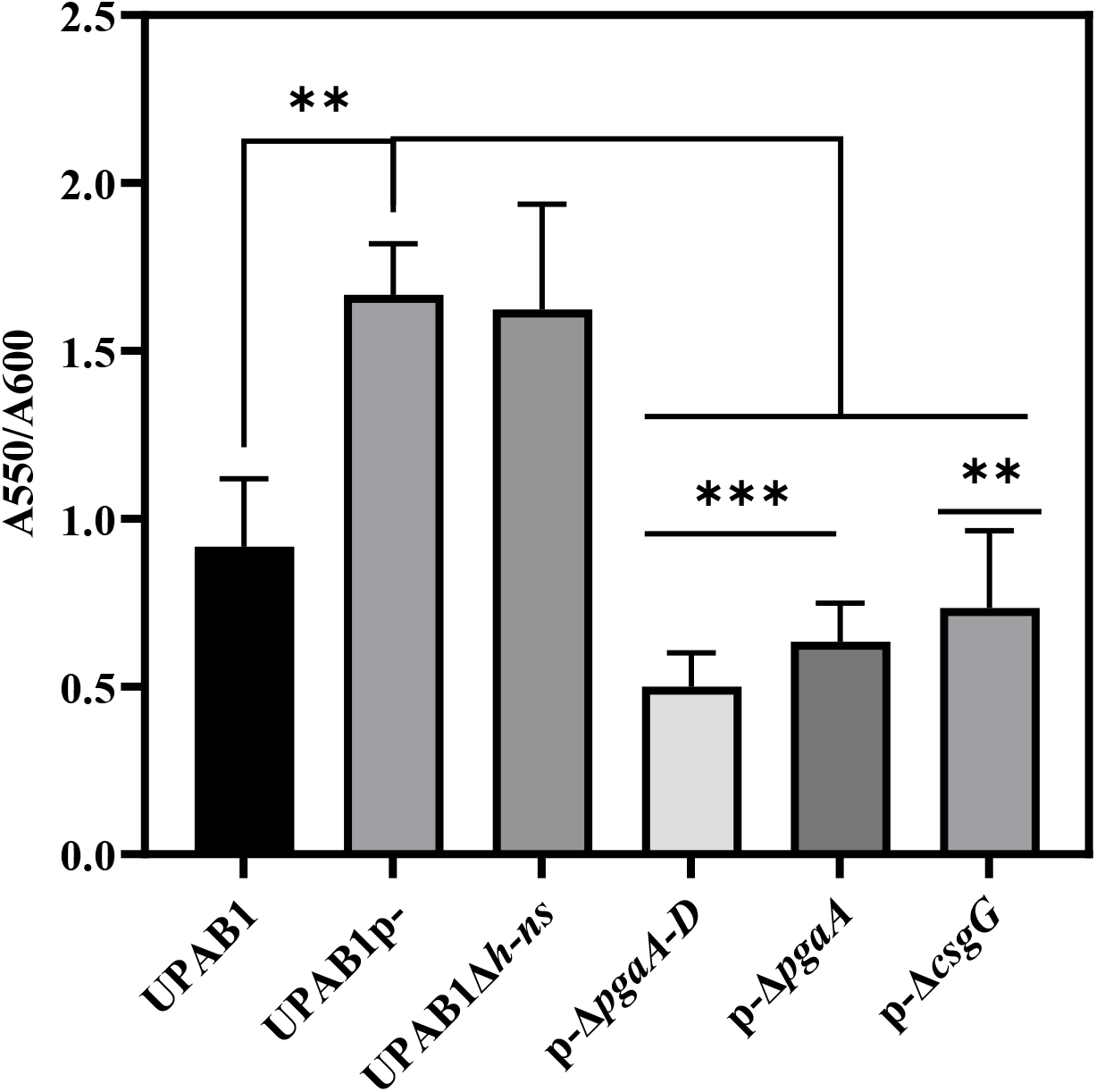
pAB5 reduce biofilm formation. Cells were grown for 8 hours on LB broth at 37°C in static conditions. Biofilm formation was measured by crystal violet and normalized to the growth. The values represent the mean and standard deviations from three independent experiments. T-test was performed by comparison with the pAB5-strain (^***^ p≤0.0005, ^**^ p≤ 0.005)

### The *csgG* cluster is widely distributed in Gram-negative bacteria

Our results show that CsgG is implicated in different phenotypes in UPAB1. A bioinformatics analysis revealed that this cluster is widespread among Gram-negative bacteria (fig 9). In some bacterial species, such as *Neisseria meningitidis* or *Vibrio harveyi*, the locus contains additional genes predicted to be co-transcribed. Recently, the crystal structure of GNA1162 from *N. meningitidis*, a homologue to D1G37_12605, has been solved (21). GNA 1162 exhibited structural similarities to TolB, and authors speculate that this protein may act as an accessory protein to an unidentified transport machinery. Despite, this cluster is present in a very large number of bacteria, its roles remain unknown and further studies are needed to identify the target of this cluster and its importance in virulence.

**Figure 9.**
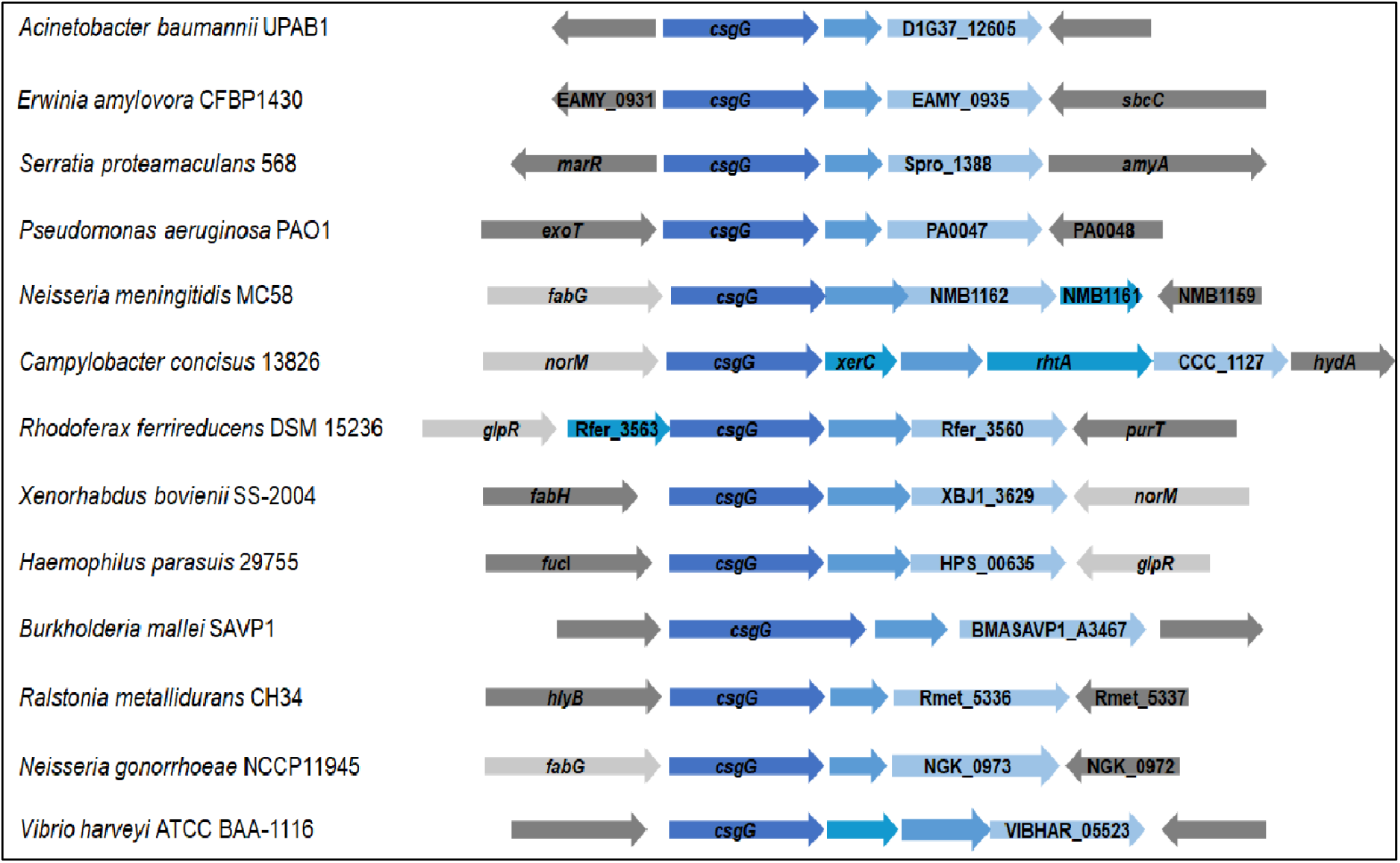
Distribution of *csgG* cluster in Gram-negative bacteria. The *csgG* cluster is indicated in blue.

## DISCUSSION

We have previously shown that the *Acinetobacter* LCPs play a key role in the dissemination of MDR and influences the pathogenesis of this bacterium by controlling the expression of chromosomally-encoded virulence factors. In this study, we show that pAB5, the LCP from UPAB1, encodes a H-NS transcriptional regulator that inhibits biofilm formation by repressing the expression of PNAG. Additionally, we found that pAB5-encoded HN-S represses the expression of a three-gene cluster that contains a homolog of CsgG, a protein involved in curli assembly. We found that disruption of CsgG dramatically reduced PNAG production, CUP pili assembly, and consequently, biofilm formation. Bioinformatic analysis revealed that the “curli like cluster” is widely distributed among Gram-negative bacteria without any attributed function.

The cross-regulatory pathways between plasmids and the bacterial chromosomes have been recently reviewed (22). Plasmid-encoded transcriptional regulators can modulate the expression of genes involved in many different processes, including motility, glycogen synthesis, adherence and quorum sensing, among others (22). H-NS-like proteins are encoded in plasmids in *Shigella flexneri* and *Salmonella enterica*. However, in these species, plasmid-encoded H-NS appear to regulate only genes horizontally acquired which maintain the energetic cost of their expression at a lower level, without affecting expression of other chromosomal genes (22). We have previously shown that repressing chromosomally encoded T6SS enable the dissemination of LCPs via conjugation (10). By downregulating PNAG, CsgG, and CUP pili, pAB5 represses biofilm formation in UPAB1. In monospecies biofilms, bacteria are surrounded by their kin, which limits the dissemination capacity of the plasmids. It is tempting to speculate that promoting the planktonic lifestyle of the host increases the chances of dissemination of pAB5 to other bacterial hosts. However, further work is required to understand the physiological implication of this process.

Our discovery that pAB5-encoded H-NS represses biofilm formation in UPAB1 is preceded by similar findings in *E. coli* and *Salmonella* (23-28). However, in these species H-NS is encoded in the chromosome. In *E. coli* and *Salmonella*, the CsgG protein is part of a multi protein complex responsible for curli fiber formation, and H-NS regulates its expression by repressing *csgD*, a key regulator for curli synthesis. In these species, H-NS is part of an intricate regulatory network that integrates diverse environmental conditions and ultimately controls curli biogenesis. CsgG is part of a predicted operon together with two putative lipoproteins of unknown function that is widely distributed in Gram-negative bacteria. *Acinetobacter* spp does not encode a complete curli biosynthetic machinery and curli fibers have not been reported. In *Caulobacter*, the CsgG ortholog, named HfaB is part of a cluster containing HfaABD, where HfaA has properties of amyloid proteins (CsgA). The HfaABD complex is critical for anchoring holdfast, a polysaccharide made of N-acetyl-d-glucosamine (NAG), and other sugars to the *Caulobacter* cell surface. It has been proposed that holdfast is attached to HfaA by an unknown mechanism (29-31). We hypothesize that a similar function anchoring PNAG to the cell surface is accomplished by a multimeric complex formed by CsgG and the two lipoproteins in the operon. Further work is necessary to determine the exact role of this gene cluster in the assembly of the extracellular matrix in these species.

## MATERIALS AND METHODS

### Bacterial strains and growth conditions

Bacterial strains, plasmids, and oligonucleotides used in this study are listed in Supplementary information. Unless otherwise noted, all strains were grown in Lysogeny broth (LB) broth at 37C with shaking (200 rpm). For strain constructions, we used gentamicin (15 or 20 μg ml^-1^), kanamycin (7.5 or 50 μg ml^-1^), zeocin (50 μg ml^-1^ with low salt LB), hygromycin (300 μg ml^-1^).

### Construction of *A. baumannii* mutants and complement strains, and pVRL2 constructs

Plasmids, and oligonucleotides used in this study are listed in Supplementary Tables S2 and S3, respectively. The constructs for generating deletions in *h-ns, pgaA-D, pgaA, csgG* and D1G37_126000 were made by substitution of the gene by an antibiotic (kanamycin or zeocin) cassette as described previously (32). Selection of mutants was carried out using the proper antibiotic. To make unmarked strains, electrocompetent mutants were transformed with pAT03 to remove the FRT-flanked antibiotic cassette. Transformants were plated on LB-agar plates containing 2 mM IPTG + hygromycin to express FLP recombinase. All strains were verified by antibiotic resistance, PCR amplification and gene sequencing. To generate genetic complementation, genes of interest were cloned into the pUC18T-miniTn7T-Gm (zeo) vector and introduced to UPAB1p-strains via four-parental mating methods as described previously (33,34). Briefly, 100 l of stationary cultures was normalized to an OD600 of 2.0 of each recipient strain, and HB101(pRK2013), EC100D(pTNS2), and EC100D containing the pUC18T-miniTn7T constructs were added to 600 μl of warm LB. Each suspension was washed twice by centrifugation at 7000g, followed by resuspension of the bacterial pellet in 1 ml of warm LB. On the final wash, the bacterial pellet was resuspended in 25 μl of LB, and the suspension was spotted on a prewarmed LB agar plate (or low-salt LB agar plate) and incubated overnight at 37°C. The bacteria were scraped from the plate, resuspended in 1 ml of LB, vortexed, and serial dilutions were plated on L agar plates supplemented with chloramphenicol to select against *E. coli* strains and gentamicin or zeocin to select for *A. baumannii* strains that had received the mini-Tn7 constructs. Correct insertion of the constructs was verified by PCR amplification and sequencing. The expression of different regulators of the pAB5 plasmid as p/*h-ns*, p/*tetR*1 and others were made using the pVRL2 vector (13). Constructs were generated by restriction enzime cloning using the HindIII and PstI sites. All the constructs were introduced to UPAB1p-strains by electroporation and transformants were selected on gentamicin.

### Congo red plate and congo red binding quantification

The red versus white color of cells was investigated using YESCA agar media (35) supplemented with 50 μg/ml Congo red. To quantify the congo red binding for bacteria prestained on the YESCA congo red plates (36). Cells were recovered from YESCA congo red plates after incubation at 26C for 48 hours. Cells were washed twice in 50 mM potassium phosphate buffer by centrifugation at 16,000 x g for 2 min and resuspended in 1 ml 50 mM potassium phosphate buffer, and the OD was adjusted to 1.100 µl of each sample were loaded onto a 96-well opaque plate, the fluorescence of congo red was measured using the plate reader (BioTek microplate spectrophotometer) with an excitation wavelength at 485nm and emission at 612 nm. The buffer was used as the blank.

### Reverse transcription-PCR

Cells were taken from LB plates after an overnight growth at 26°C and normalized to OD 1 and treated with RNA protect. RNA purification was prepared using the Quick RNA fungal/bacterial miniprep (Zymo Research) by following the manufacturer’s instructions with some modification in the DNA digestion step as follows. Contaminating DNA was removed using the Turbo DNA-free kit (Invitrogen) by following the manufacturer’s instructions. For reverse transcription (RT)-PCR, cDNA was prepared from 1 μg RNA using a high-capacity RNA-to-cDNA kit (Applied Biosystems), according to the manufacturer’s protocol. Real-time quantitative PCR (qPCR) was performed using Power SYBR green PCR master mix reagents (Applied Biosystems) on a ViiA7 real-time PCR system (Applied Biosystems), following the manufacturer’s suggested protocol. In all cases a no-template control was run with no detectable transcripts.

### Biofilm formation

Cells were grown overnight in 5 ml of LB broth (or YESCA), then cultures were diluted to an OD600 of 0.01 in LB broth (or YESCA). Cultures were deposited in 96-well plates and incubated at 37 °C for 8 or 24 hours without shaking. Cultures were removed to read the absorbance at 600nm. Then, plates were washed three times with water, stained with 0.1% Crystal Violet (w/v) and quantified at 550 nm after solubilization with 30% acetic acid.

### Detection of PNAG production

Immunoblotting to detect PNAG production was performed as described previously (14) with some modification. Cells grown overnight on LB, then diluted to an OD of 1 and spotted in to YESCA plate and incubated for 48 hours at 26 °C. Cells were scraped from plate and normalized to an OD of 1, then pelleted and resuspended in 300 µl of 0.5 M EDTA (pH 8.0). Cells were incubated for 5 min at 98 °C, and centrifuged at 9000 x g for 5 min. After centrifugation, supernatants were diluted 1:3 in Tris-buffered saline (TBS) and incubated with 100 µl of proteinase K (20 mg/ml) for 60 min at 65 °C then for 30 min at 80 °C (to inactivate the protease). The preparations were serially diluted in TBS, and 5 µl were spotted on nitrocellulose membrane and let the membrane dry completely. Next, membrane was blocked and incubated with an anti-PNAG antibody (kind gift of Dr. Gerald B. Pier, Harvard Medical School), and an anti-human IgG (IRDye 800 CW) antibody (LI-COR Biosciences, Lincoln, NE) and visualized with an Odyssey CLx imaging system (LI-COR Biosciences). Following, membrane was incubated with anti-UPAB1 primary antibody (Ref) followed by an incubation with anti-rabbit IgG (IRDye 800 CW) antibody (LI-COR Biosciences, Lincoln, NE).

### Scanning Electron Microscopy

Overnight cultures on YESCA media were diluted in YESCA + 4% DMSO (37) to an OD600 of 0.02 in 24 well-plate containing glass coverslips and incubated at 26°C for 24 hours with shaking. Then, the media was removed, and the 24 well-plate was washed with 0.15 M cacodylate buffer. Cells were fixed overnight at room temperature on a shaker using the fixative solution (2.5% glutaraldehyde, 2% paraformaldehyde and 0.2% tannic acid in 0.15M cacodylate buffer pH 7.4 with 2mM calcium chloride). Post fixation, coverslips were rinsed in 0.15 M cacodylate buffer 3 times for 10 minutes each followed by a secondary fixation in 1% OsO4 in 0.15 M cacodylate buffer for 45 minutes in the dark. The coverslips were then rinsed 3 times in ultrapure water for 10 minutes each and dehydrated in a graded ethanol series (10%, 30%, 50%, 70%, 90%, 100% x2) for 10 minutes each step. Once dehydrated, the samples were loaded into a critical point drier (Leica EM CPD 300, Vienna, Austria) which was set to perform 12 CO_2_exchanges at the slowest speed. Once dried, coverslips were mounted on aluminum stubs with carbon adhesive tabs and coated with 10 nm of carbon and 6 nm of iridium (Leica ACE 600, Vienna, Austria). SEM images were acquired on a FE-SEM (Zeiss Merlin, Oberkochen, Germany) at 1.5 kV and 0.1 nA.

### Transmission electron microscopy and cup pili detection

Overnight cultures on YESCA media were diluted in YESCA+ 4% DMSO (37) to an OD600 of 0.02 and incubated at 26°C for 48 hours with shaking. Next cultures were washed with PBS and used for TEM and western blot. For negative staining and analysis by transmission electron microscopy, Bacterial samples were fixed with 1% glutaraldehyde (Ted Pella Inc., Redding CA) and allowed to absorb onto freshly glow discharged formvar/carbon-coated copper grids for 10 min. Grids were then washed in dH_2_O and stained with 1% aqueous uranyl acetate (Ted Pella Inc.) for 1 min. Excess liquid was gently wicked off and grids were allowed to air dry. Samples were viewed on a JEOL 1200EX transmission electron microscope (JEOL USA, Peabody, MA) equipped with an AMT 8-megapixel digital camera (Advanced Microscopy Techniques, Woburn, MA). For cup pili detection, cells were resuspended in Laemmli buffer to a final OD of 10. Samples were loaded onto 15% SDS-PAGE gel for separation, transferred to a nitro-cellulose membrane and probed with polyclonal rabbit anti CupA (1:1000) and monoclonal mouse anti-RNA polymerase (1:3000, Biolegend, San 397 Diego, CA). Western blots were then probed with IRDye-conjugated anti-mouse and anti-rabbit secondary antibodies (both at 1:15,000, LI-COR Biosciences, Lincoln, NE) and visualized with an Odyssey CLx imaging system (LI-COR Biosciences).

### Generation of polyclonal rabbit sera against CupA

The UPAB1 gene *cupA* was cloned into pET28a+ with a 10-histidine tag using primers SB cupA bamh1 F and SB cupA hind3 R, creating pET28-CupA10His, and electroporated into *E. coli* DH5α. pET28-CupA10His was confirmed by sequencing. E. coli Rosetta 2 cells were used for CupA purification. 1liter of LB was inoculated from an overnight culture of Rosetta 2/pET28-CupA10His at an OD_600_ of 0.05. Culture grown to an OD_600_ of ∼ 0.5 before induction with 1 mM isopropyl 1-thio-β-d-galactopyranoside (IPTG). The cultures were grown for an additional 4 hours. Cells were harvested at 12000 x g for 20 min. Cells were washed with cold PBS and resuspended in binding buffer supplemented with protease inhibitor (300 mM NaCl, 10 mM imidazole, 30 mM Tris-HCl, pH 8.0). Cells were lysed with a cell disruptor using three rounds at 35 kp.s.i (Constant System Ltd., Kennesaw, GA). Cell lysates were centrifuged at 20000g (or 11000 rpm) for 20 min to collect inclusion bodies. Pellet was resuspended in binding buffer and centrifuged as described before twice. Then pellet was resuspended in binding buffer containing urea and incubated for 3 hours at 4°C with continuous stirring (6 M urea, 300 mM NaCl, 10 mM imidazole, 30 mM Tris-HCl, pH 8.0). Then, lysates were centrifuged at 35000 x g for 20 min and supernatant was filtered using 0.45 µm filter. Next, Cell lysates were passed over a nickel-NTA agarose column (Gold Bio, St. Louis, MO) equilibrated with 10 column volumes of binding buffer. The load fraction is the total cell lysate. The flow-through was collected as what passed through the column and did not bind the nickel-NTA resin. The column was washed first with 15 column volumes of washing buffer (5M urea, 20 mM imidazole, 300 mM NaCl, 30 mM Tris-HCl, pH 8.0) and second with 10 column volumes of washing buffer (4M urea, 20 mM imidazole, 300 mM NaCl, 30 mM Tris-HCl, pH 8.0). Proteins were eluted using elution buffer (2M urea, 250 mM imidazole, 300 mM NaCl, 30 mM Tris-HCl, pH 8.0). Elution fractions were analyzed by SDS-PAGE analysis and Coomassie staining. The polyacrylamide gel band corresponding to CupA-His was sent to Antibody Research Corporation (St. Louis, MO) for peptide extraction and development of rabbit-derived polyclonal antibodies.

### Growth assays

Bacteria were cultured overnight in YESCA liquid media at 26 °C under shaking conditions. Cultures were washed with PBS and diluted to an OD_600_ of 0.01 in 150 µL of YESCA liquid media in 96 well plates and incubated at 26 °C under shaking conditions. OD_600_ values were measured every 30 min for 16 hours via a BioTek microplate spectrophotometer. Three separate experiments were performed with four wells per experiment for each strain.

## ACKNOWLEDGEMENTS

We acknowledge the assistance of Dr. Wandy Beatty at the Molecular Microbiology department Imaging Facility at Washington University School of Medicine in transmission electron microscopy studies, Dr. Sanja Sviben and Dr. James Fitzpatrick at the Washington University Center for Cellular Imaging (WUCCI) in scanning electron microscopy studies, which is supported by Washington University School of Medicine, The Children’s Discovery Institute of University and St. Louis Children’s Hospital (CDI-CORE-2015-505 and CDI-CORE-2019-813), the Foundation for Barnes-Jewish Hospital (3770). We thank Dr. Gerald Pier for sharing the anti-PNAG antibody.

This work was supported by grants from the National Institute of Allergy and Infectious Diseases (grants R01AI144120 and R01AI125363).

## SUPPLEMENTAL MATERIAL

**Figure S1.**
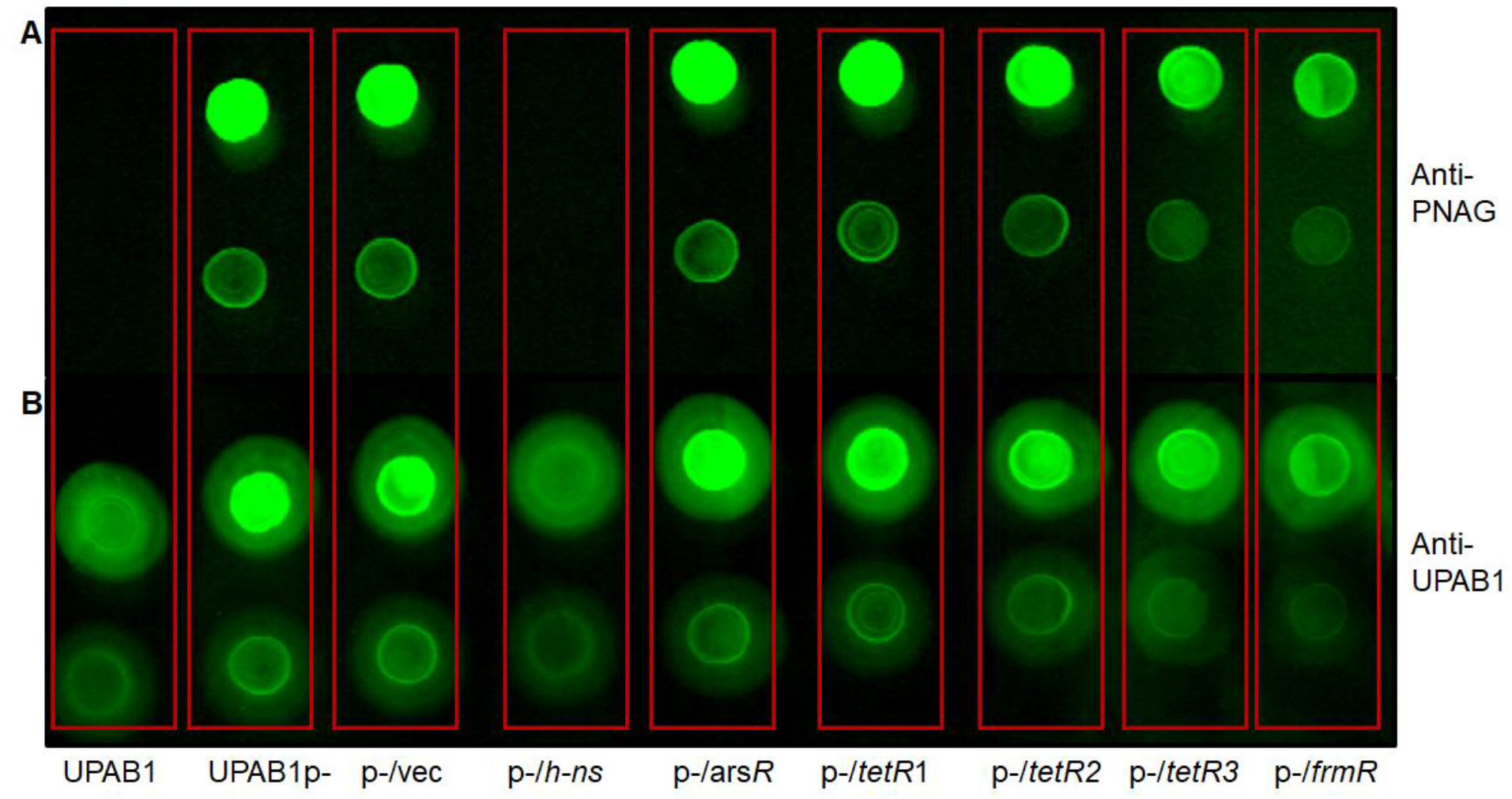
H-NS reduce PNAG production. UPAB1, UPAB1p-, and UPAB1p-empty vector (p-/vec) or the vector expressing the pAB5 regulators. Immunoblots using antibodies anti-PNAG (A) and antibodies anti-UPAB1 as loading control (B). Cells were taken from overnight LB-agar plates incubated at 26 °C and adjusted to OD 1.

**Figure S2.**
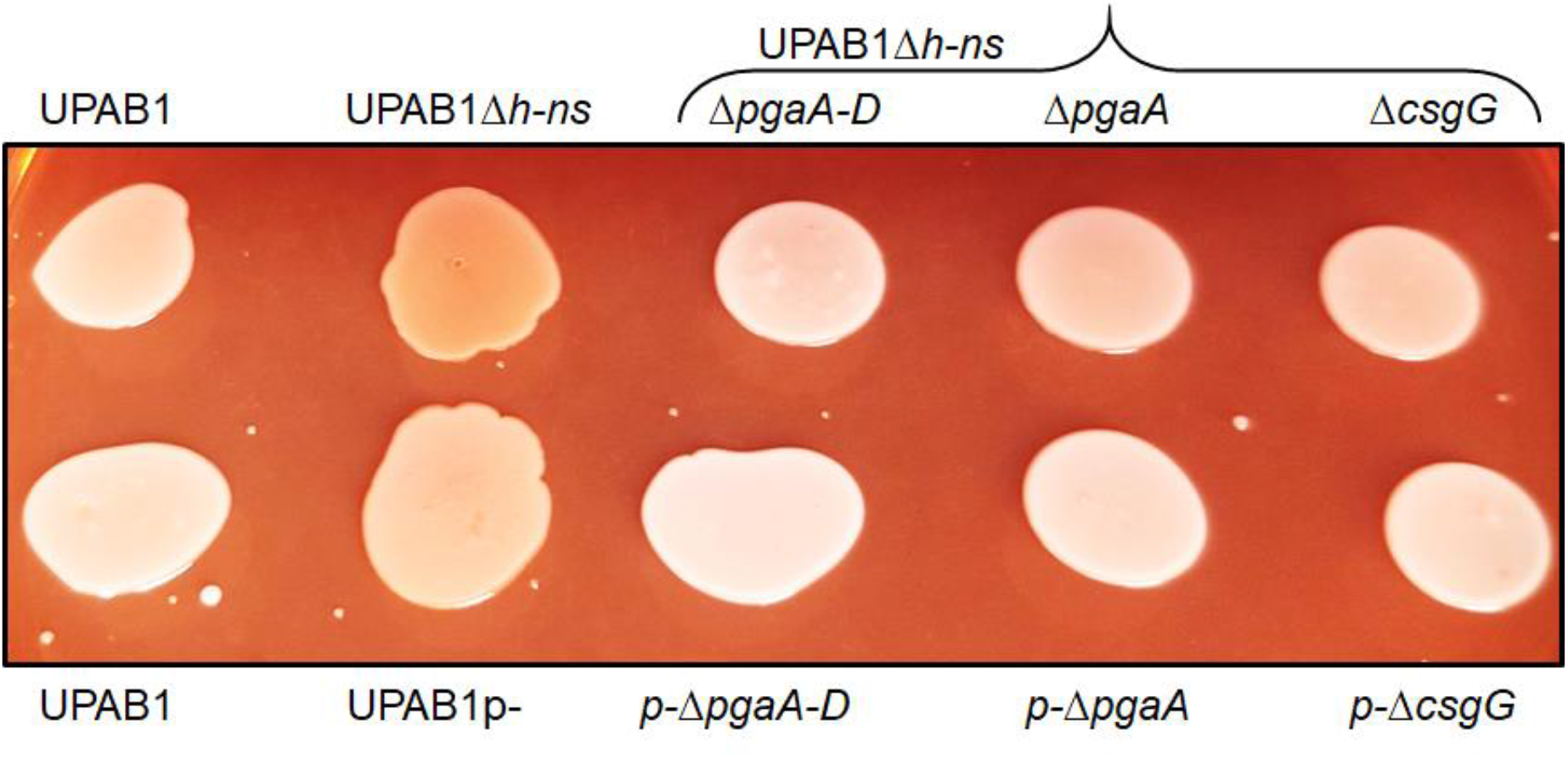
PgaA-D and CsgG like clusters are involved in Congo-red binding. Deletions of pgaA-D, pgaA and csgG were made in UPAB1Δ*h-ns* background (top panel) and UPAB1p-background. Cells were incubated for 48 hours at 26°C on YESCA-Congo-red agar plates.

**Figure S3.**
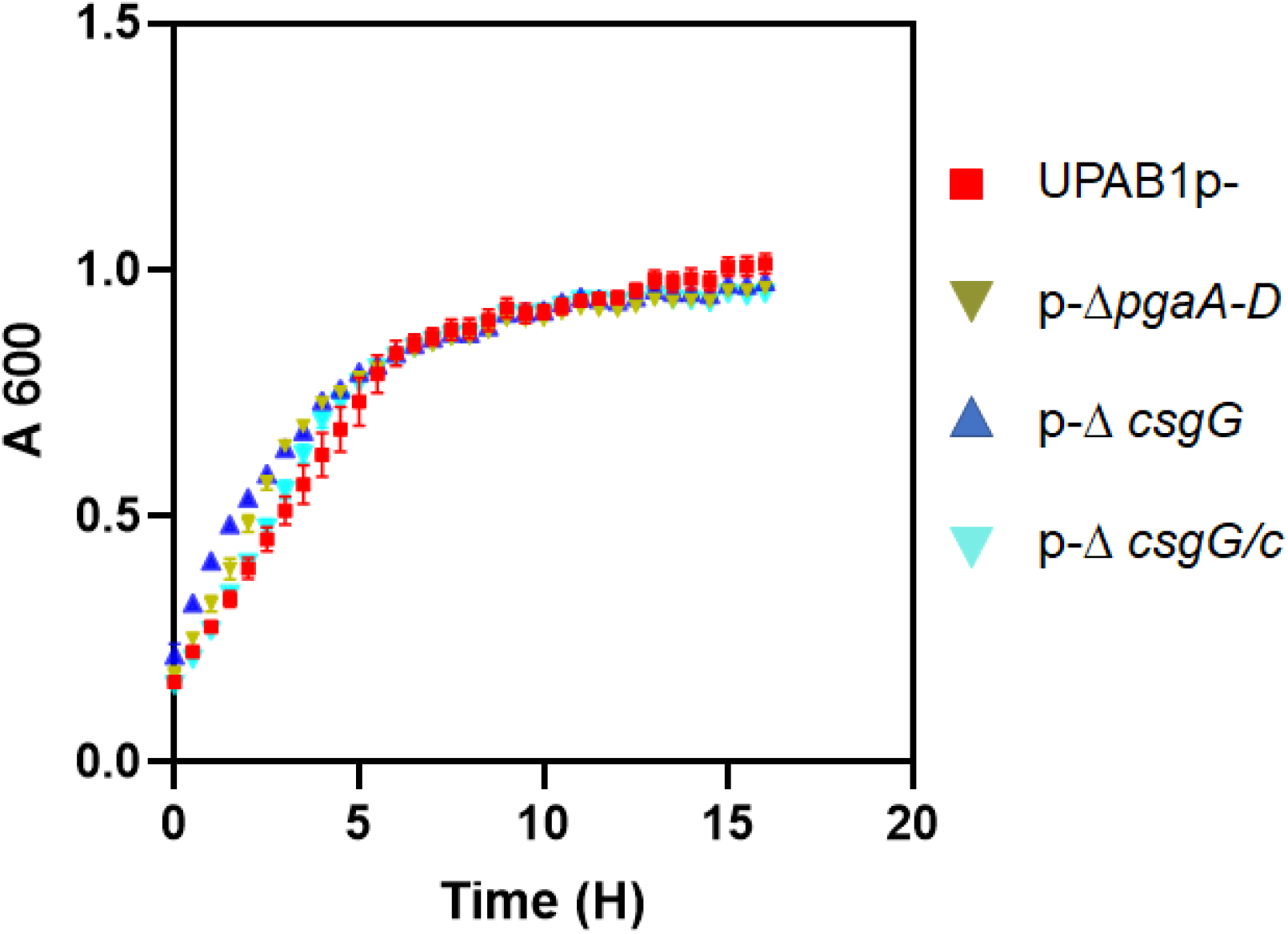
Growth curves of UPAB1p- and derivative mutant strains in YESCA-DMSO media, measured by OD600. The graphs represent the mean and standard deviation of three replicates.

**Figure S4.**
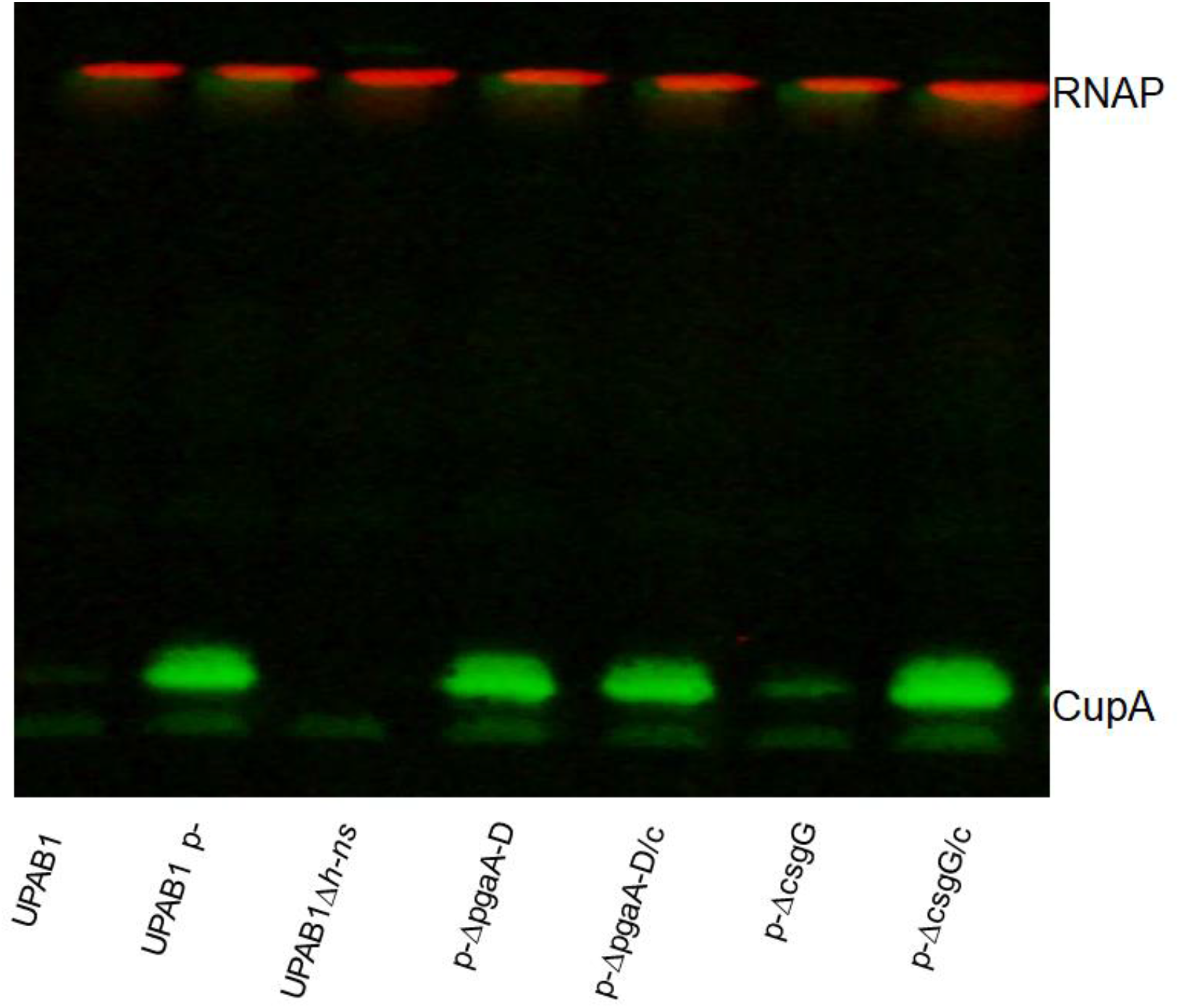
pAB5 inhibits CUP pili formation. Western blot of OD-normalized whole cell. Western blot probing for CupA (the Usher protein from Cup 2 pili). RNAP is included as loading control.

**Figure S5.**
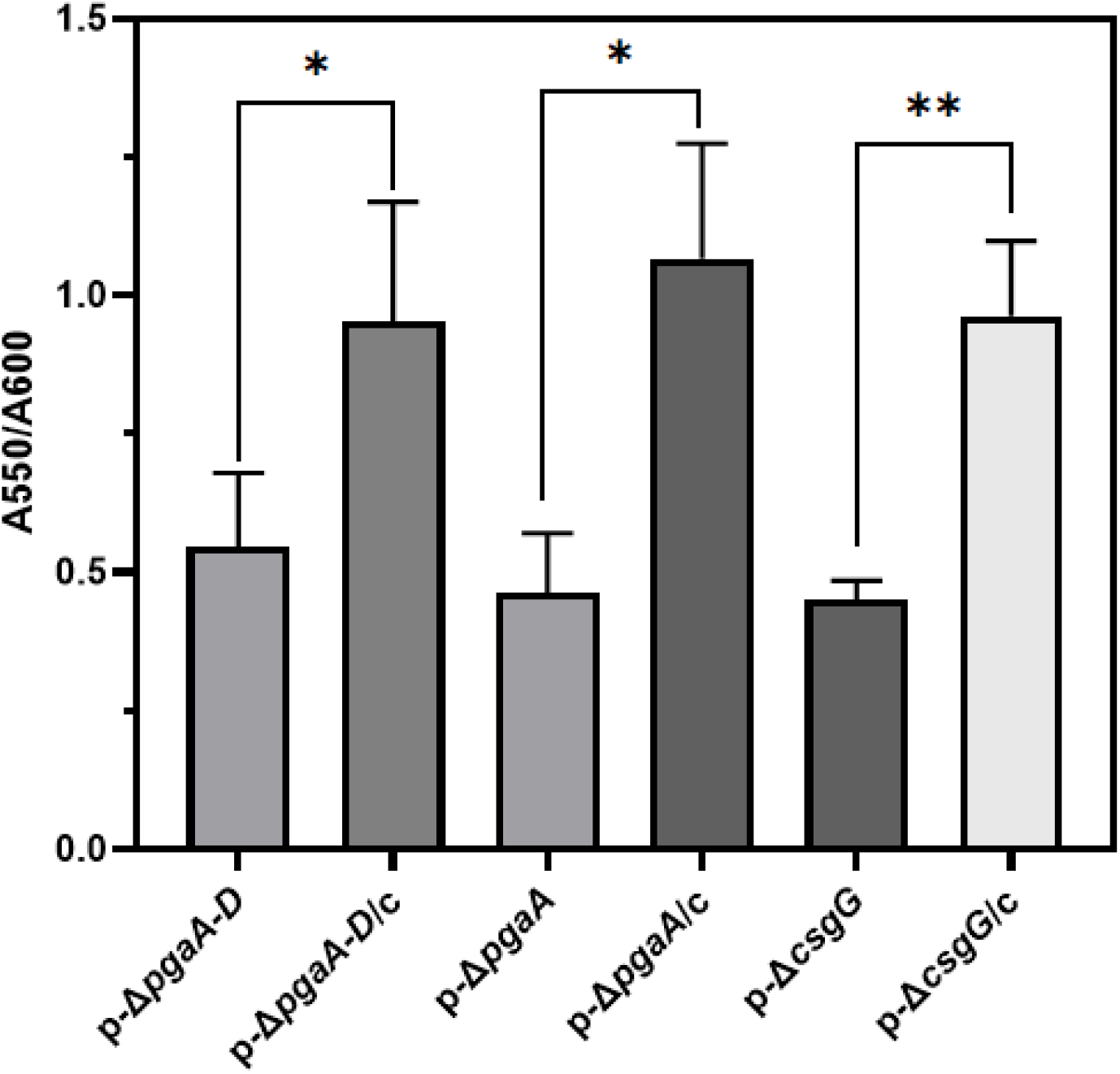
Biofilm formation in UPAB1p-mutants and complemented strains. Cells were grown for 8 hours on LB broth at 37°C under static conditions. Biofilm formation was measured by the crystal violet binding and normalized to the OD600. The values represent the mean and standard deviations from three independent experiments. Statistical analysis by t test was performed by comparison with the pAB5-strain (^**^ p≤0.005, ^*^ p≤ 0.05).

**Table S1.**
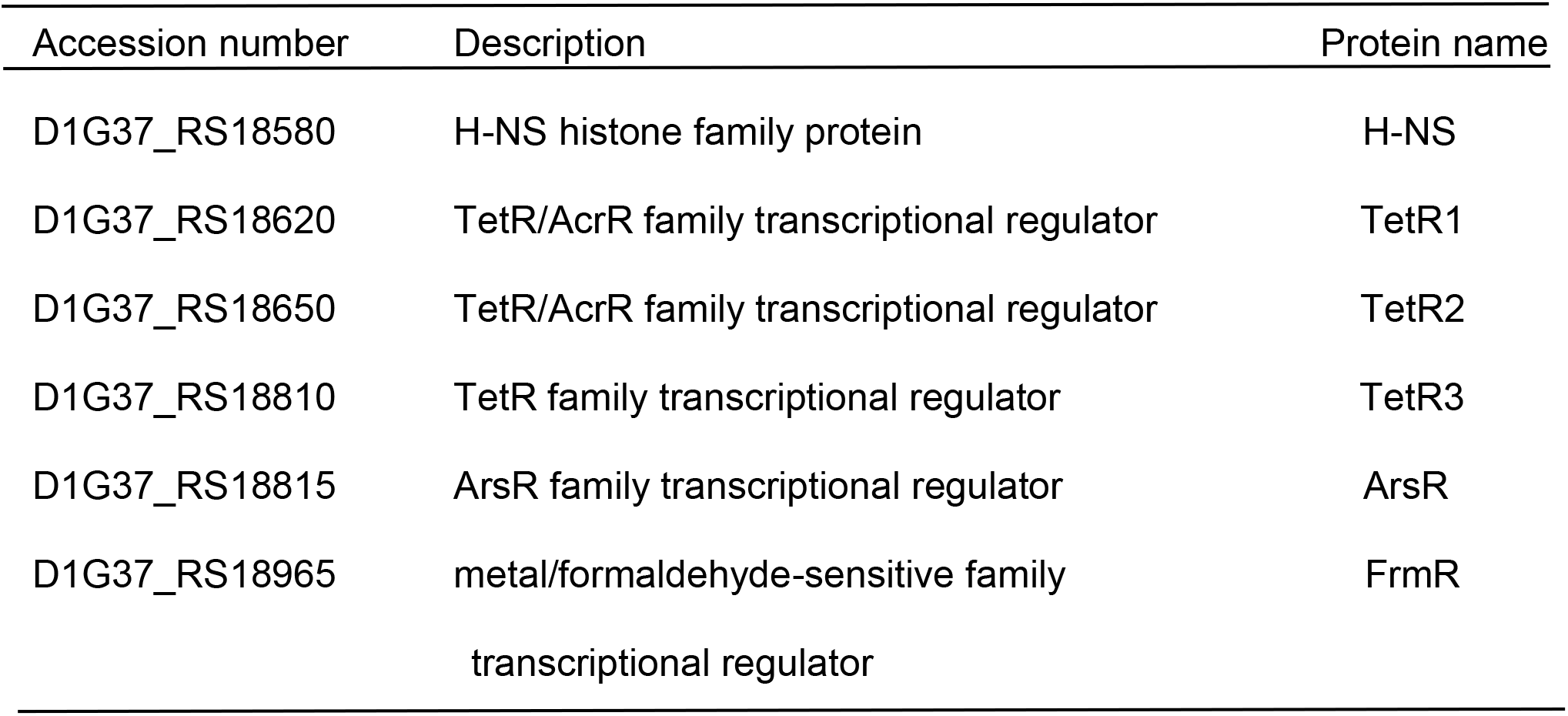
List of regulators identified in pAB5.

| 716 | Accession number | Description | Protein name |
| --- | --- | --- | --- |
| 717 | D1G37_RS18580 | H-NS histone family protein | H-NS |
| 718 | D1G37_RS18620 | TetR/AcrR family transcriptional regulator | TetR1 |
| 719 | D1G37_RS18650 | TetR/AcrR family transcriptional regulator | TetR2 |
| 720 | D1G37_RS18810 | TetR family transcriptional regulator | TetR3 |
| 721 | D1G37_RS18815 | ArsR family transcriptional regulator | ArsR |
| 722 | D1G37_RS18965 | metal/formaldehyde-sensitive family | FrmR |
| 723 |  | transcriptional regulator |  |

**Table S2.** Bacterial strains used in this study.

| 739 | Strain | Relevant properties | Reference |
| --- | --- | --- | --- |
| 740 | UPAB1 | MDR Urine isolate with pAB5 plasmid | (1) |
| 741 | UPAB1p- (p-) | UPAB1 derivative strain without pAB5 | (1) |
| 742 | UPAB1$\Delta h$-ns | UPAB1 containing an unmarked *h*-ns deletion | this study |
| 743 | p-/vec | UPAB1p- containing the pVRL2 expression vector | this study |
| 744 | p-/h-ns | p- expressing *h*-ns in pVRL2 | this study |
| 745 | p-/tetR1 | p- expressing *tetR1* in pVRL2 | this study |
| 746 | p-/tetR2 | p- expressing *tetR2* in pVRL2 | this study |
| 747 | p-/tetR3 | p- expressing *tetR3* in pVRL2 | this study |
| 748 | p-/arsR | p- expressing *arsR* in pVRL2 | this study |
| 749 | p-/frmR | p- expressing *frmR* in pVRL2 | this study |
| 750 | p-$\Delta pgaA$-D | p- containing an unmarked *pgaA*-D deletion | this study |
| 751 | p-$\Delta pgaA$ | p- containing an unmarked *pgaA* deletion | this study |
| 752 | p-$\Delta csgG$ | p- containing an unmarked *csgG* deletion | this study |
| 753 | p-$\Delta Ab$ 12600 | p- containing an unmarked D1G37_12600 deletion | this study |
| 754 | p+$\Delta h$-ns $\Delta pgaA$-D | UPAB1 containing an unmarked *h*-ns | this study |
| 755 |  | and *pgaA*-D deletion |  |
| 756 | p+$\Delta h$-ns $\Delta pgaA$ | UPAB1 containing an unmarked *h*-ns | this study |
| 757 |  | and *pgaA* deletion |  |
| 758 | p+$\Delta h$-ns $\Delta csgG$ | UPAB1 containing an unmarked *h*-ns | this study |
| 759 |  | and *csgG* deletion |  |
| 760 | p-$\Delta pgaA$-D/c | p-$\Delta pgaA$-D complemented | this study |
| 761 | p-$\Delta pgaA$/c | p-$\Delta pgaA$ complemented | this study |
| 762 | p-$\Delta csgG$/c | p-$\Delta csgG$ complemented | this study |

**Table S3.**
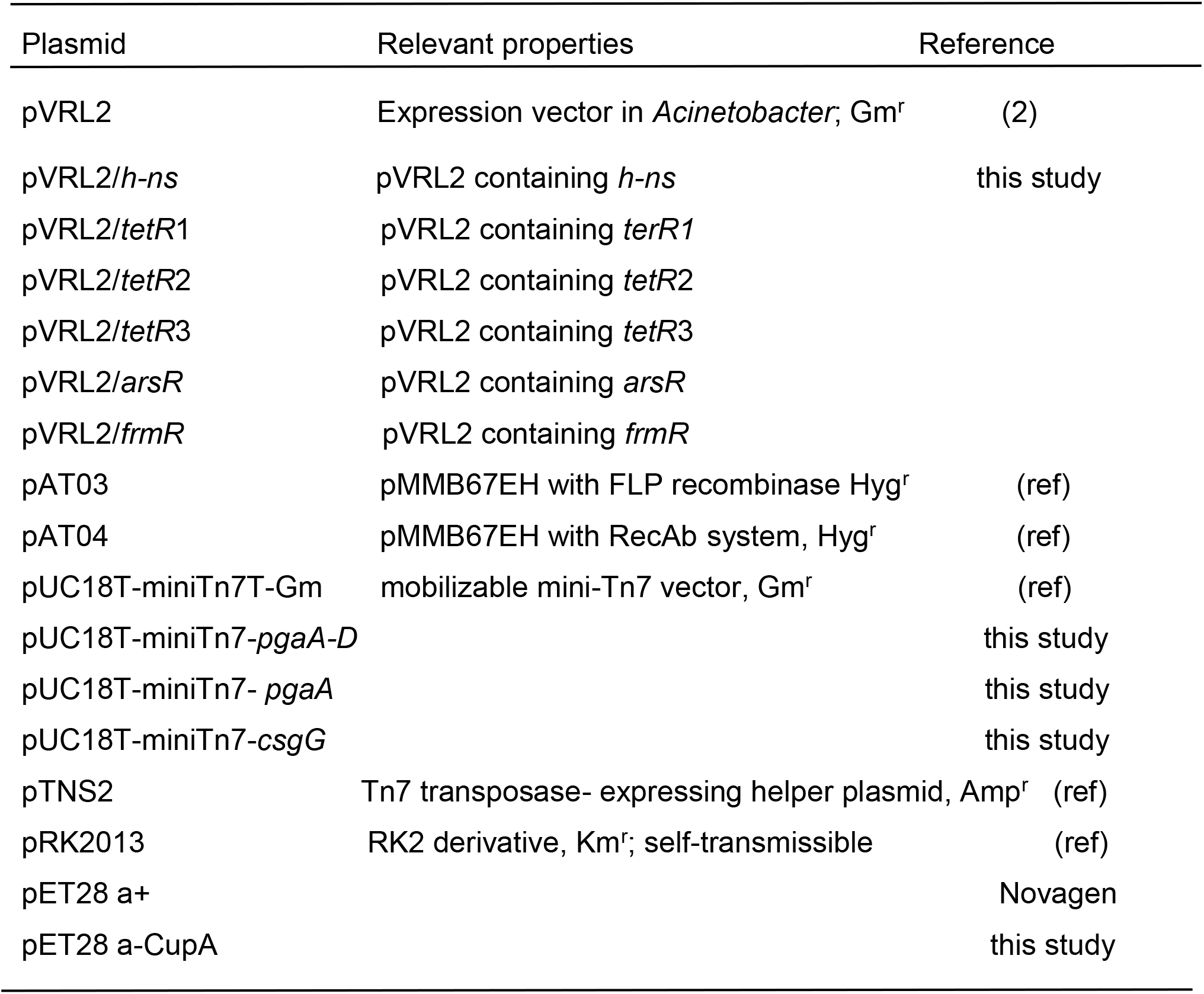
Bacterial plasmids used in this study.

